# Activation of STING due to COPI-deficiency

**DOI:** 10.1101/2020.07.09.194399

**Authors:** Annemarie Steiner, Katja Hrovat Schaale, Ignazia Prigione, Dominic De Nardo, Laura F. Dagley, Chien-Hsiung Yu, Pawat Laohamonthonkul, Cassandra R. Harapas, Michael P. Gantier, Marco Gattorno, Stefano Volpi, Sophia Davidson, Seth L. Masters

**Affiliations:** Inflammation Division, The Walter and Eliza Hall Institute of Medical Research, Parkville, VIC 3052, Australia; Department of Medical Biology, University of Melbourne, VIC 3010, Australia; Centro per le Malattie Autoinfiammatorie e Immunodeficienze, IRCCS Istituto Giannina Gaslini, Genoa, Italy; Department of Anatomy and Developmental Biology, Monash Biomedicine Discovery Institute, Monash University, Clayton, Victoria 3168, Australia; Advanced Technology and Biology, The Walter and Eliza Hall Institute of Medical Research, Parkville, VIC 3052, Australia; Centre for Innate Immunity and Infectious Diseases, Hudson Institute of Medical Research, Clayton, VIC 3168, Australia; Department of Molecular and Translational Science, Monash University, Clayton, VIC 3168, Australia; Universita’ degli Studi di Genova, Genoa, Italy

**Keywords:** COPA syndrome, inflammation, Stimulator of interferon genes (STING), type I interferons, retrograde trafficking

## Abstract

COPA syndrome is caused by loss-of-function mutations in the COP-α subunit of coatomer protein complex I (COPI), which participates in retrograde vesicular trafficking of proteins from the Golgi to the endoplasmic reticulum (ER). Disease manifests early in life with arthritis, lung pathology, kidney dysfunction and systemic inflammation associated with NF-κB activation and type I interferon (IFNαβ) production. Here, we generated *in vitro* models for COPA syndrome and interrogated inflammatory signalling pathways via a range of biochemical and molecular biological techniques. Results were confirmed with cell lines in which mutant COPA was overexpressed and with COPA syndrome patient PBMCs. We identified Stimulator of Interferon Genes (STING), as a driver of inflammation in COPA syndrome. Furthermore, we found that genetic deletion of COPG1, another COPI subunit protein, induced NF-κB and type I IFN pathways similar to COPA-deficiency. Finally, we demonstrate that *in vitro*, inflammation due to COPA syndrome mutations was ameliorated by treatment with the small molecule STING inhibitor H-151. Therefore, inflammation induced by deletion of COPI subunits in general suggests a link between retrograde trafficking and STING regulation, and this innate immune sensor represents a novel therapeutic target in COPA syndrome.

## Introduction

Coatomer subunit α (COPA) syndrome is a recently identified rare disorder, involving complex pathology with dysregulation of the innate and adaptive immune system (1). The disease is caused by autosomal dominant mutations in the *COPA* gene, which encodes the α–subunit of the coatomer complex I (COPI). COPI mediates retrograde trafficking of cargo proteins from Golgi to endoplasmic reticulum (ER) and within cis-Golgi compartments (2). To date, 11 families with mutations in *COPA* have been identified worldwide (1, 3-10). Clinically, patients present symptoms with different severity, including interstitial lung disease with or without pulmonary haemorrhage, inflammatory arthritis and immune-mediated kidney disease. Autoantibodies and increased levels of Th17 cells have been identified in the majority of patients (1).

Most striking is the disease onset during childhood or early adulthood and the incomplete penetrance, leaving some individuals unaffected, despite carrying the mutation (1). Interestingly, across all reported COPA syndrome patients, a total of 5 missense mutations within exon 8 and 9 of the *COPA* gene have been identified, which translates into a 14 amino acid stretch within the WD40 domain of the COPA (COPα) protein. Being highly conserved between species and implicated in protein-protein interactions, COPA mutants were shown to have impaired binding efficiency to cargo proteins, therefore causing defective retrograde transport (1). As a consequence, ER stress, activation of the unfolded protein response (UPR) and nuclear factor kappa B (NF-κB) pathway activation were suggested pathomechanisms, which have already been linked to lung disease and autoimmunity in other studies (11-13). Furthermore, Volpi and colleagues described elevated transcription levels of type I interferons (IFNs) and interferon-stimulated genes (ISGs) in peripheral blood of COPA syndrome patients, suggesting a role of type I IFN signalling and a dysregulated innate immune response in disease pathogenesis (3). This finding is supported by two recent case reports showing therapeutic benefit of JAK1/2 inhibition in COPA syndrome patients (10, 14). However, the innate immune sensor as well as the molecular mechanisms underlying COPA syndrome pathogenesis remain unclear.

Due to the involvement of COPA in intracellular trafficking and its subcellular localization, we wondered about a role of COPA in STING (Stimulator of IFN gene, also known as MITA, MPYS and ERIS) (15-18) pathway regulation. Inactive STING forms homodimers that localize to the ER membrane and has been identified as an adapter protein downstream of multiple intracellular nucleotide sensors including cyclic-GMP-AMP-synthase (cGAS) (19, 20). Upon recognition of cytosolic double stranded DNA as a danger-associated molecular pattern (DAMP), cGAS synthesizes 2’3’-cGAMP, a second messenger detected by STING (21). Secondary messenger binding results in a conformational change in STING, which enables translocation to the Golgi apparatus. This is an essential step for downstream signalling to occur (22). At the Golgi compartment, transcription factors such as NF-κB and IFN regulatory factor 3 (IRF3) are activated via IκB kinase epsilon (IKKε) and TANK-binding-kinase 1 (TBK1) in a phosphorylation-dependent manner (23). Translocation of these transcription factors to the nucleus then results in gene expression of proinflammatory cytokines and type I and III IFNs (IFNαβ and IFNλ) (24, 25). Identifying the innate immune pathway activated in COPA syndrome will provide valuable insights in disease pathology leading to beneficial pharmacological intervention for this inflammatory condition.

## Methods

### Cell culture

All used cell lines were obtained from ATCC. Human embryonic kidney (HEK293T) cells and HeLa cells were cultured in DMEM (Gibco) supplemented with 100U/ml Penicillin (Sigma-Aldrich),100μg/ml Streptomycin (Sigma-Aldrich) and 10% Fetal Bovine Serum (FBS, Sigma Aldrich) in a humidified incubator at 37°C and 10% CO_2_. The human monocytic THP-1 cell line was maintained in RPMI-1640 (made inhouse, RPMI 1640 powder (Life Technologies), 23.8mM Sodium Bicarbonate (NaHCO_3_) (Merck), 1mM Sodium Pyruvate (C_3_H_3_NaO_3_) (Sigma-Aldrich), 100U/ml Penicillin, 100μg/ml Streptomycin) supplemented with 10% FBS and incubated in humidified atmosphere at 37°C and 5% CO_2_.

### CRISPR/Cas9 gene editing

COPA^deficient^, MAVS^-/-^, PKR^-/-^, UNC93B1^-/-^, NLRP3^-/-^, STING^-/-^ cells were generated using CRISPR/Cas9 gene editing as previously described (26). In brief, lentiviral transduction was performed to stably express the FU-Cas9-mCherry plasmid (Addgene #70182) or lentiCRISPR V2 Cas9 (28, 29) in THP-1 or HeLa cells. Therefore, 3rd generation lentiviral particles were generated by transient transfection of packaging vectors pMDL (5µg), VSVg (3µg), Rsv-Rev (2.5µg) and plasmid of interest (10µg) into one 10 cm dish of HEK293T cells. After 24 hours, media was changed to 10 ml complete growth medium. 24 hours later, 3 ml of lentivirus-containing supernatant was used to transduce 0.5-1×10^6^ THP-1 cells or 0.2-0.5×10^6^ HeLa cells by spin infecting for 3 hours at 3200 rpm and 32°C in presence of polybrene (4µg/ml). Stable cell lines were selected by flow cytometry associated cell sorting (FACS) or puromycin selection (5µg/ml, Invivogen), respectively. Single guide (sg) RNAs (sequences listed in Suppl. Table 1) were cloned into the Doxycycline (Dox)-inducible FgH1t-UTG or FgH1t-UTC construct (Addgene #70183), followed by lentiviral transduction of Cas9-expressing THP-1 and HeLa cell lines. Pools of stable cell lines were selected by FACS.

Gene deletion in pooled cell lines was induced by treatment with Dox (1 μg/ml, Sigma Aldrich) for 48-72 hours prior to each experiment. Protein or gene deletion was assessed by western blotting or quantitative Real-Time PCR (qRT-PCR). Dox-inducible double knockout cell lines were generated by deletion of signalling molecules MAVS, PKR, STING, NLRP3 and UNC93B1 in THP-1 or HeLa cells stably expressing Cas9 and COPA-specific sgRNA1. Stably Cas9-expressing THP-1or HeLa cells were used as control cells and are referred to as parental cell line throughout this study.

### Immunoblotting

0.5 – 1×10^6^ cells were lysed in RIPA buffer (20mM Tris-HCl (pH 7.3), 150mM NaCl, 5mM EDTA, 1% Triton X-100, 0.5% Sodium Deoxycholate, 0.1%SDS, 10%Glycerol) supplemented with cOmplete™ Protease Inhibitor Cocktail (11697498001, Roche Biochemicals, Mannheim, Germany), 1mM PMSF, 5mM NaF, 10mM NaPPi, 1mM Na_3_VO_4_) at 1×10^6^ cells/100µl. Using Pierce centrifuge columns (Thermo Fisher Scientific), protein lysates were purified of DNA and subsequently mixed with SDS-PAGE sample buffer. Size-based protein separation was achieved by use of 4-12% SDS PAGE gels (Novex) and MES-running buffer (Thermo Fisher Scientific). Proteins were then transferred onto a PVDF membrane (Immobilon^®^-P Transfer Membrane, Millipore) via wet transfer and blocked in 5% skim milk/Tris-buffered saline (TBST) for 1 hour at RT. Membranes were probed with primary antibody diluted in 5% skim milk/TBST or 5% Bovine Serum Albumin (BSA)/TBST over night at 4°C: anti-COPA (Santa Cruz Biotechnology, clone H-3, sc-398099), anti-phospho-STAT1 Tyr701 (Cell Signaling Technology, clone 58D6, #9167), anti-phospho-TBK1 Ser172 (Cell Signaling Technology, clone D52C2, #5483), anti-phospho-IRF3 Ser386 (Abcam, ab76493), anti-STING (Cell Signaling Technology, clone D2P2F, #13647), anti-STAT1 (Cell Signaling Technology, clone D1K9Y, #14994), anti-TBK1 (Cell Signaling Technology, #3031), anti-COPG (A-10, sc-393977, Santa Cruz Biotechnolgy), anti-SEC13 (F-3, sc-514308, Santa Cruz Biotechnology) or anti-Actin-HRP (Santa Cruz Biotechnology, clone C4, sc-47778). Secondary antibody was incubated for 1 hour at room temperature (1:10000, sheep anti-mouse IgG-HRP or donkey anti-rabbit IgG-HRP, GE Healthcare). For development, Immobilon Forte Western HRP Substrate (Cat. No. WBLUF0500, Millipore) and the ChemiDoc Imaging touch system (BioRad) was used.

### RNA isolation and quantitative Real-Time PCR

RNA was isolated using the ISOLATE II RNA Mini Kit (Bioline) following the manufacturer’s guidelines. The Superscript III Reverse transcriptase (Invitrogen) and oligo (dT) nucleotides (Promega) were used to reverse transcribe 1μg of total RNA. Quantitative Real-Time PCR (q-RT-PCR) was performed using SYBR Green/ROX qPCR Master Mix (Thermo Fisher Scientific) and the ViiA 7 Real-time PCR system (Thermo Fisher Scientific). Primers used in this study are listed in Suppl. Table 2. Samples were run in duplicates, normalized to housekeeper gene *ACTIN* and analysed using the ΔΔCt method. Data are presented as fold change relative to vehicle control or parental cell line.

### Inhibitors

STING inhibitor studies were performed during 72 hours Dox treatment. Therefore, THP-1 cells were seeded at 8×10^4^ cells/6-well plate in 2ml complete RPMI medium (+Dox 1μg/ml). After 34 hours incubation, STING inhibitor H-151 (2.5μM, Invivogen) or DMSO (vehicle control) was added and allowed to incubate for 11 hours. Cells were washed and incubation with Dox continued until 72 hours time point was reached.

### Transient transfection

The pCMV6-Entry-COPA-myc-DDK plasmid (Origene) was used as a template to generate COPA mutant E241K and R233H expressing plasmids via QuikChange Lightning Multi Site-Directed Mutagenesis (AgilentTechnologies) according to the manufacturer’s instructions. For overexpression studies, HEK293T cells were initially seeded at 2×10^5^ cells/well in a 6 well plate format and transiently transfected with 0.5μg of total DNA as stated in the figure or figure legend. Transfection of all plasmids including pEF-BOS-mCitrine-hSTING and pEF-BOS-FLAG as an empty vector (EV) control (kindly provided by V.Hornung, LMU Munich), was performed with Lipofectamine 2000 (Thermo Fisher Scientific) reagent following the manufacturer’s instructions. After 24 hours incubation, cells were lysed for immunoblot analysis.

### Immunoprecipitation of mCitrine-STING for mass spectrometry

Immunoprecipitation (IP) experiments were performed similarly as described (27). Briefly, 3×10^6^ HEK293T cells were seeded in 10cm dishes. The following morning cells were transiently transfected with 10µg of either pEF-BOS EV control or pEF-BOS-mCitrine-hSTING plasmid DNA. After 48 hours cells were lysed in 750µl 1xNonidet P-40 (NP-40) [1% Nonidet P-40, 10% Glycerol, 20mM Tris-HCl pH 7.4, 150mM NaCl, 1mM EGTA, 10mM NaPPi, 5mM NaF, 1mM, Na_3_VO_4_] supplemented with 1mM PMSF and 1 x cOmplete protease inhibitors (Roche Biochemicals). WCLs were clarified by centrifugation at 17,000 x g for 10min at 4°C. For each IP sample, 1µg of mouse 1gG2A monoclonal GFP antibody (Thermo Fisher Scientific, [E36] A-11120) was crosslinked to 50μl of Protein A Dynabeads→ (Thermo Fisher Scientific, 10002D) using 5mM BS^3^ (Thermo Fisher Scientific) and incubated for 30min at RT. The crosslinking reaction was quenched by adding 1M Tris-HCl [pH 7.4]. Antibody-crosslinked Protein A Dynabeads→ were thoroughly washed before 50μl was then added to each IP sample and incubated on a rotator at 4°C for 2 hours. Beads were then washed four times with 1xNP-40 using a DynaMag-2 magnetic holder (Thermo Fisher Scientific) before proteins were eluted with 0.5% SDS in PBS. Eluted protein material was subjected to tryptic digestion using the FASP method as previously described (28-30). Peptides were lyophilised using CentriVap (Labconco) prior to reconstituting in 60μl 0.1% FA/2% acetonitrile. Peptide mixtures (2μl) were separated by reverse-phase chromatography on a C18 fused silica column (I.D. 75 µm, O.D. 360 μm x 25 cm length) packed into an emitter tip (IonOpticks), using a nano-flow HPLC (M-class, Waters). The HPLC was coupled to an Impact II UHR-QqTOF mass spectrometer (Bruker) using a CaptiveSpray source and nanoBooster at 0.20 Bar using acetonitrile. Peptides were loaded directly onto the column at a constant flow rate of 400nl/min with 0.1% formic acid in MilliQ water and eluted with a 90 min linear gradient from 2 to 34% buffer B (99.9% acetonitrile and 0.1% formic acid). Mass spectra were acquired in a data-dependent manner including an automatic switch between MS and MS/MS scans using a 1.5 second duty cycle and 4Hz MS1 spectra rate, followed by MS/MS scans at 8-20Hz dependent on precursor intensity for the remainder of the cycle. MS spectra were acquired between a mass range of 200–2000m/z. Peptide fragmentation was performed using collision-induced dissociation. Raw files consisting of high-resolution MS/MS spectra were processed with MaxQuant (version 1.6.6.0) for feature detection and protein identification using the Andromeda search engine (31) as previously described (32). Extracted peak lists were searched against the reviewed *Homo sapiens* (UniProt, March 2019) database as well as a separate reverse decoy database to empirically assess the false discovery rate (FDR) using strict trypsin specificity, allowing up to 2 missed cleavages. LFQ quantification was selected, with a minimum ratio count of 2. PSM and protein identifications were filtered using a target-decoy approach at an FDR of 1%. Only unique and razor peptides were considered for quantification with intensity values present in at least 2 out of 3 replicates per group. Statistical analyses were performed using LFQAnalyst (33) (https://bioinformatics.erc.monash.edu/apps/LFQ-Analyst/) whereby the LFQ intensity values were used for protein quantification. Missing values were replaced by values drawn from a normal distribution of 1.8 standard deviations and a width of 0.3 for each sample (Perseus-type). Protein-wise linear models combined with empirical Bayes statistics were used for differential expression analysis using Bioconductor package Limma whereby the adjusted *p*-value cutoff was set at 0.05 and log2 fold change cutoff set at 1. The Benjamini-Hochberg (BH) method of FDR correction was used.

### Cytokine measurements

To measure protein levels of CXCL10 and IFNβ, THP-1 cells were seeded at 1.5×10^5^ cells per well into a 96 well plate after 48 hours of Dox treatment. 24 hours later, culture supernatants were collected and cytokine concentrations measured using CXCL10/IP10 (Human CXCL10/IP-10 Quantikine ELISA Kit, R&D Systems) and IFN-β (VeriKine-HS Human IFN Beta Serum ELISA Kit Cat No 41415, PBL Bioscience) ELISA Kits following the manufacturer’s instructions.

### Immunofluorescence

Glass coverslips (18mm x 18mm, thickness 11/2, Zeiss) were sterilised in 100% ethanol and placed in 6 well plates. After 72 hours of Dox treatment, HeLa cells were seeded at 3×10^5^ cells/well in 2ml complete growth medium and left to attach for 6-8 hours. Following treatment with HT-DNA (2μg/ml, transfected with Lipofectamine 2000 (Thermo Fisher Scientific)) for 2 hours, cells were washed in 1xPBS and fixed in 2ml ice cold methanol for 15min at -20°C. Prior to incubation in blocking buffer (5% normal goat serum, 0.3% Triton-×100, 1xPBS) at RT for 1 hour, cells were washed 3 times in 1xPBS. Primary antibodies were diluted in antibody dilution buffer (1% BSA, 0.3% Triton-×100, 1xPBS) and incubated overnight at 4°C using the following dilutions: anti-COPA (1:100, Santa Cruz Biotechnology, clone H-3, sc-398099) and anti-STING (1:100, Cell Signaling Technology, clone D2P2F, #13647). Next, cover slips were washed 3 times in 1xPBS and incubated with secondary antibodies (anti-rabbit-AF647, anti-mouse-AF488 (BD Bioscience)) diluted 1:1000 in antibody diluent for 1 hour at RT. Cover slips were washed 3x in 1xPBS and mounted onto microscopy slides using Fluoromount-G with DAPI (Thermo Fisher Scientific). Z-stack images were acquired using Zeiss LSM 880 NLO Microscope and analysed with Fiji software. Microscopical images are presented as maximum intensity projections.

### Patient PBMCs and FACS analysis

Human blood samples were collected after informed consent (ethical review board of Instituto Gianina Gaslini-Genolva-Italy N. BIOL 6/5/04). PBMCs from a COPA syndrome patient, previously reported in (3), and two healthy control (HC) donors were treated with STING inhibitor H-151 (5μM, InvivoGen) at 37°C for 4 hours. Cells were fixed with pre-warmed Fixation Buffer (BioLegend) at 37° for 15 min and permeabilized with pre-chilled True-Phos(tm) Perm Buffer (BioLegend) at -20° C for 1 hour. Cells were then stained with CD3-APC mAb (SK7, BD Biosciences), CD14-FITC mAb (M5E2, BD Biosciences) and phospho-TBK1/NAK (Ser172) (D52C2, Cell Signaling Technology), XP Rabbit mAb-PE (Cell Signaling Technology) or Rabbit (DA1E) mAb IgG XP Isotype Control-PE (Cell Signaling Technology) at RT for 30 min. Samples were analysed using the FACSCanto A (BD Biosciences) flow cytometer and FlowJo 10.5 software. The monocyte population was firstly identified based on cell size and granularity and subsequently confirmed by gating for CD14-positive/CD3-negative subpopulation. Data for phosphorylated TBK1 (pTBK1) is presented as histogram showing mean fluorescence intensity (MFI) for the CD14-expressing monocyte population. Quantification (Fig. 3B) shows fold change of MFI (representing pTBK1) after H-151 treatment and was calculated using this formula:

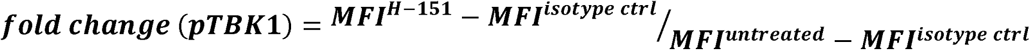

### Statistical analysis

Unless stated otherwise in the figure legends, data are presented as mean of biological replicates (n=3). Error bars represent standard error of the mean (SEM). Using the GraphPad Prism8 software, statistical comparison was made either by paired ratio t-test or one-way ANOVA as stated in the figure legends (p-value < 0.05 considered statistically significant).

## Results

### Generation of cellular models for COPA syndrome

Aiming to study the protein function of COPA in the context of innate immune signalling we used CRISPR/Cas9 genome editing to delete COPA in human monocytic THP-1 and epithelial HeLa cell lines. This approach was based on the assumption that the reduced target protein binding efficiency of loss of function mutations identified in COPA syndrome patients can be mimicked by reduction of wildtype COPA protein levels. We generated three COPA^deficient^ THP-1 pooled cell lines using different single guide (sg) RNAs targeting different exons of the *COPA* gene. In the monocytic human THP-1 cell line, reduction of COPA protein levels coincided with spontaneous phosphorylation of the transcription factor signal transducer and activator of transcription 1 (STAT1), which is downstream of type I and III IFN signalling (Fig. 1A). Complete deletion of COPA could not be achieved because it is essential for cell survival (34). We proceeded to use the THP-1 cell line with sgRNA 1 (termed COPA^deficient^) as it demonstrated the greatest reduction in COPA levels and concomitant increase in STAT1 phosphorylation (pSTAT1). An elevation of proinflammatory cytokine gene expression levels (*IL1B, IL6, IL12, IL4, IL23*) has previously been described in B cell lines derived from COPA syndrome patients and a type I IFN signature was reported in patient PBMCs (1, 3). Similarly, qRT-PCR analysis of our COPA^deficient^ THP-1 cells revealed increased mRNA expression levels of proinflammatory cytokines (*TNF* and *IL6*), the type I IFN subtypes, *IFNA1, IFNB1*, as well as interferon-stimulated genes (ISGs), such as *ISG15, IFIT1* and *MX1* (Fig. 1B). Furthermore, increased levels of IFNβ and the IFN-induced chemokine, CXCL10 proteins were measured by ELISA in supernatants from COPA^deficient^ THP-1 cells (Fig. 1C).

**Figure 1:**
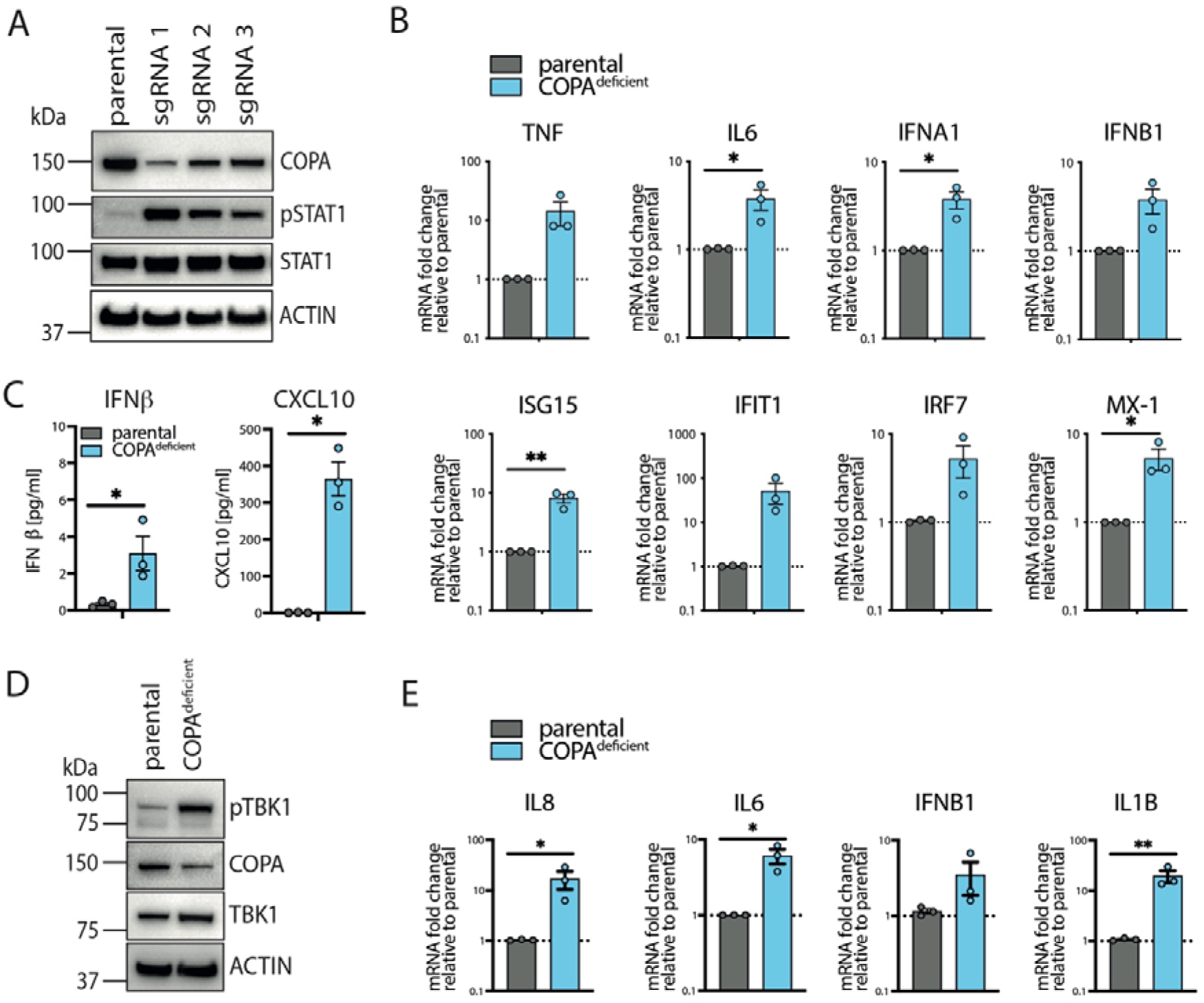
Cell line models of COPA syndrome. **A)** CRISPR/Cas9 gene editing technology was employed to generate pooled COPA^deficient^ THP-1 cell lines. Three different single guide (sg) RNAs targeting exon 5 (sgRNA1), exon1 (sgRNA2) or exon7 (sgRNA3) of the COPA gene were used. Baseline protein levels of COPA, STAT1 and phosphorylated STAT1 (pSTAT1) were assessed by immunoblotting, using actin as a loading control. **B)** qRT-PCR analysis of transcription levels of proinflammatory cytokines, type I IFN and ISGs at baseline was performed for a representative COPA^deficient^ THP-1 cell line (sgRNA 1). Data are pooled from 3 independent experiments and presented as fold change to parental control cell line (THP-1-Cas9). Statistical significance was assessed by ratio paired t-test, * p<0.05, ** p<0.01. **C)** Analysis of baseline levels of IFNβ and CXCL10 in cell culture supernatants by ELISA. Data are pooled from 3 independent experiments. Error bars represent SEM, statistical analysis: ratio paired t-test, * p<0.05. **D)** CRISPR/Cas9 targeting of COPA with sgRNA1 in HeLa cells resulted in reduction of COPA protein levels and spontaneous phosphorylation of TBK1 (pTBK1) at baseline as assessed by immunoblotting. **E)** Assessment of baseline gene transcription profile of COPA^deficient^ HeLa cells by qRT-PCR analysis. Data are presented as described in B).

In order to independently confirm these findings in a different cell line, sgRNA1 was used to generate COPA^deficient^ HeLa cells (Fig. 1D). Similarly, the reduction of COPA protein levels resulted in increased transcription of proinflammatory cytokines and type I IFN (Fig. 1E), as well as elevated baseline phosphorylation of TBK1 (pTBK1), a signalling molecule downstream of several pattern recognition receptors (Fig. 1D). Therefore, we have successfully generated COPA^deficient^ THP-1 and HeLa cell lines that model inflammatory manifestations observed in COPA syndrome as a useful tool for future studies to investigate the molecular mechanisms underlying this disease.

### Overactive STING pathway drives inflammatory signalling in COPA syndrome model cell lines

To identify the innate immune sensor that is driving the inflammatory response in our *in vitro* models of COPA syndrome, we genetically deleted several candidate innate immune receptors in COPA^deficient^ THP-1 cells (Suppl. Fig. 1). Using this approach, we excluded the involvement of inflammasome sensor NLRP3 (Suppl. Fig. 1A), cytoplasmic RNA sensor PKR (Suppl. Fig. 1B) and RNA-sensors RIG-I and MDA-5 by deletion of shared adaptor protein mitochondrial antiviral-signalling protein (MAVS) (Suppl. Fig. 1C). Furthermore, deletion of Unc93 homolog B1 (UNC93B1), an adaptor protein essential for endosomal TLRs and cell surface TLR5 stability and signalling (35), was not able to ameliorate the inflammatory phenotype in COPA^deficient^ THP-1 cells (Suppl. Fig. 1D).

Another candidate immune sensor is STING, which functions as adapter protein that triggers inflammation downstream of multiple cytoplasmic nucleic acid sensors including cGAS. Inactive STING localizes on the ER membrane and requires anterograde trafficking to translocate to Golgi compartments upon activation (36). Although the anterograde transport route is mediated by coatomer complex II (COPII) vesicles which COPA is not a part of, all cellular secretory pathways form a network within the cell and are tightly regulated. Therefore, we hypothesized that reduced functionality of retrograde transport might potentially interfere with STING trafficking and prolong signalling. In line with this, we identified COPA as a potential interaction partner for STING in an unbiased mass spectrometry-proteomics experiment, where we analysed protein lysates following pulldown of overexpressed mCitrine (mCit)-tagged human STING in HEK293T cells (Fig. 2A). This finding was also independently confirmed by Keskitalo and colleagues (37).

**Figure 2:**
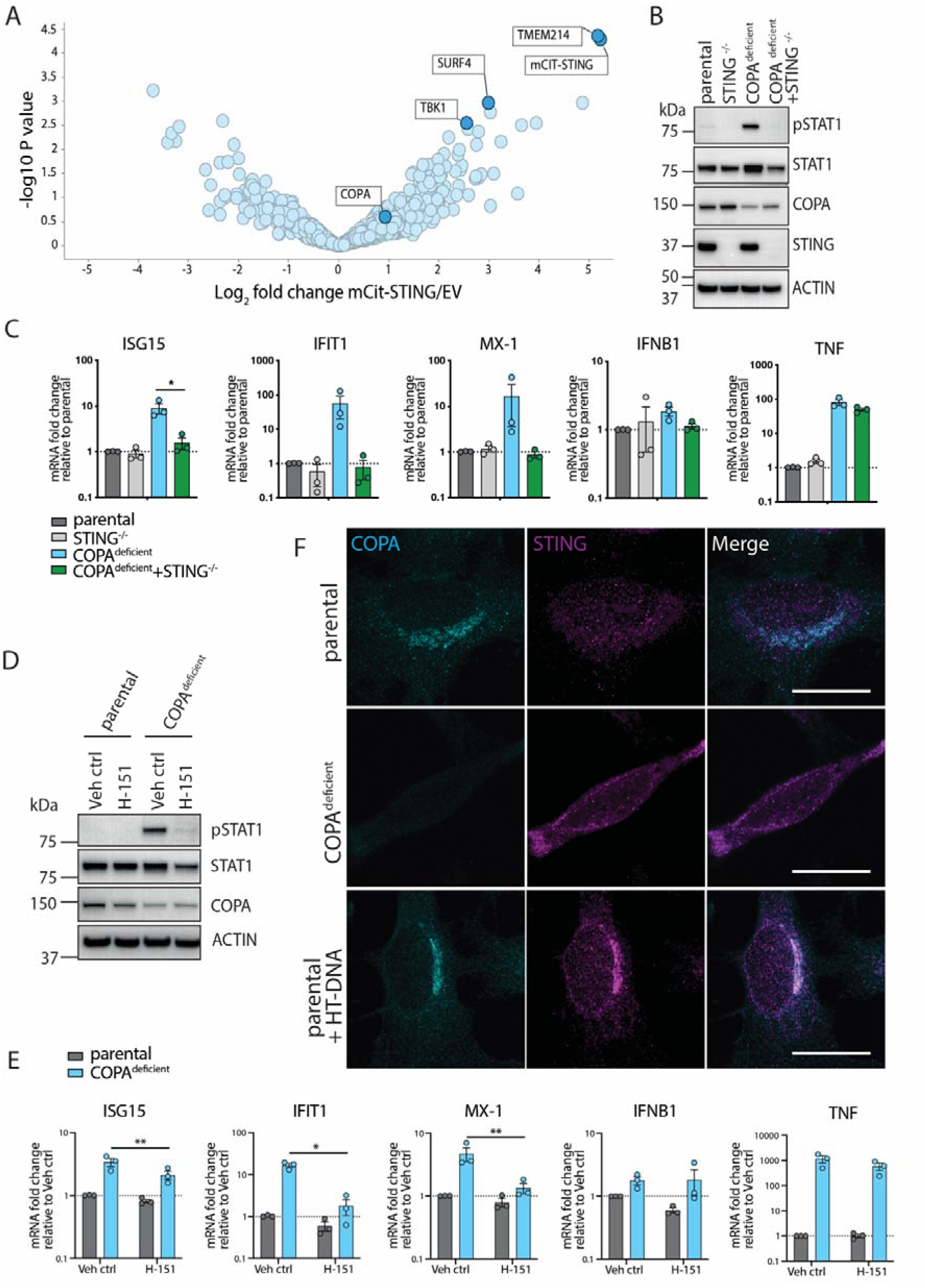
Inflammation induced by COPA-deficiency is STING dependent. **A)** Volcano plot representing the log_2_ protein ratios of differentially regulated proteins in overexpressed mCit-STING pulldowns in HEK293T cells relative to empty vector (EV) control (*n*= 3 independent IPs). COPA, other known STING interactors and TMEM214 are labelled. **B)** Using COPA^deficient^/STING^-/-^ THP-1 cells, immunoblotting was performed at baseline to assess STING, COPA and STAT1 protein levels as well as phosphorylation of STAT1 (pSTAT1). Actin was used as a loading control. **C)** qRT-PCR analysis of transcription levels in COPA^deficient^/STING^-/-^ THP-1 cells at baseline. Data are pooled from 3 independent experiments. Error bars represent SEM, statistical analysis by ratio paired t-test, * p<0.05. **D)** Immunoblotting of COPA^deficient^ THP-1 cells after treatment with STING inhibitor H-151 (2.5μM) for 11 hours was performed to assess p-STAT1 levels. Results are shown as a representative of 3 independent repeats. **E)** Analysis of proinflammatory gene and ISG transcription levels in COPA^deficient^ THP-1 cells following treatment with H-151 (2.5μM, 11 hours). Data are pooled from 3 independent experiments. Error bars represent SEM, statistical analysis: ratio paired t-test, * p<0.05. **F)** Immunofluorescence microscopy analysis was performed to study STING localization (magenta) upon deletion of COPA (cyan) in HeLa cells compared to Cas9-expressing parental control cells. HT-DNA (2μg/ml, 2 hours) treated parental HeLa cells were used as positive control showing STING activation and subsequent translocation to Golgi compartments. Representative experiment shown of n=3, scale bar 20μm.

CRISPR/Cas9-mediated genetic deletion of STING ameliorated the spontaneous pSTAT1 signal (Fig. 2B) as well as inflammatory gene expression in COPA^deficient^ THP-1 cells (Fig. 2C). Interestingly, TNF transcription levels are not significantly reduced upon STING deletion and are likely the result of NF-κB pathway activation following ER stress, as previously suggested (1). As an independent confirmation for the role of STING, genetic deletion in COPA^deficient^ HeLa cells ameliorated type I IFN-mediated inflammatory baseline signalling as assessed by qRT-PCR and immunoblot analysis for pTBK1 (Suppl. Fig. 2). In the context of potential pharmaceutical intervention, treatment of COPA^deficient^ THP-1 cells with STING inhibitor H-151 was able to ablate baseline pSTAT1 (Fig. 2D) and type I IFN-mediated gene transcription (Fig. 2E), thus indicating a possible targeted treatment for COPA syndrome patients.

Given the requirement for STING’s ER-to-Golgi translocation to activate downstream signalling molecules, we performed immunofluorescence (IF) studies and aimed to investigate whether defective retrograde transport via COPA-deficiency results in STING accumulation at the Golgi, which therefore mediates increased signalling. As positive control, parental HeLa cells were stimulated with HT-DNA, a double stranded DNA molecule to activate cGAS, which results in distinct STING translocation that can be observed as puncta formation and indicates STING activation (Fig. 2F). In COPA^deficient^ HeLa cells without further stimulation, STING did not form clear puncta like the positive control (Fig. 2F). However, considering that loss of COPA is associated with Golgi dispersal (38), localization patterns of activated STING might be significantly aberrant. Overall these results suggest that spontaneous STING activation is driving the inflammatory response in COPA syndrome model THP-1 and HeLa cell lines, although this may involve smaller signalling complexes at dispersed Golgi fragments.

### Mutations in COPA drive STING-dependent inflammation

In order to validate our findings in patient samples, we analysed PBMCs from a COPA syndrome patient carrying the c.698G>A (p.R233H) mutation with clinical presentation of severe polyarticular arthritis and lung disease (3). At the time of sample collection, the patient was treated with Prednisone and Rituximab. Flow cytometry analysis revealed elevated levels of pTBK1, particularly in CD14-expressing monocytes (Suppl. Fig. 3). Treatment with STING inhibitor H-151 was able to ameliorate baseline pTBK1 as shown in representative histograms and quantified as fold change of pTBK1 levels after H-151 treatment (measured by mean fluorescence intensity, (MFI)), thereby confirming basal STING activation (Fig. 3A, B).

**Figure 3:**
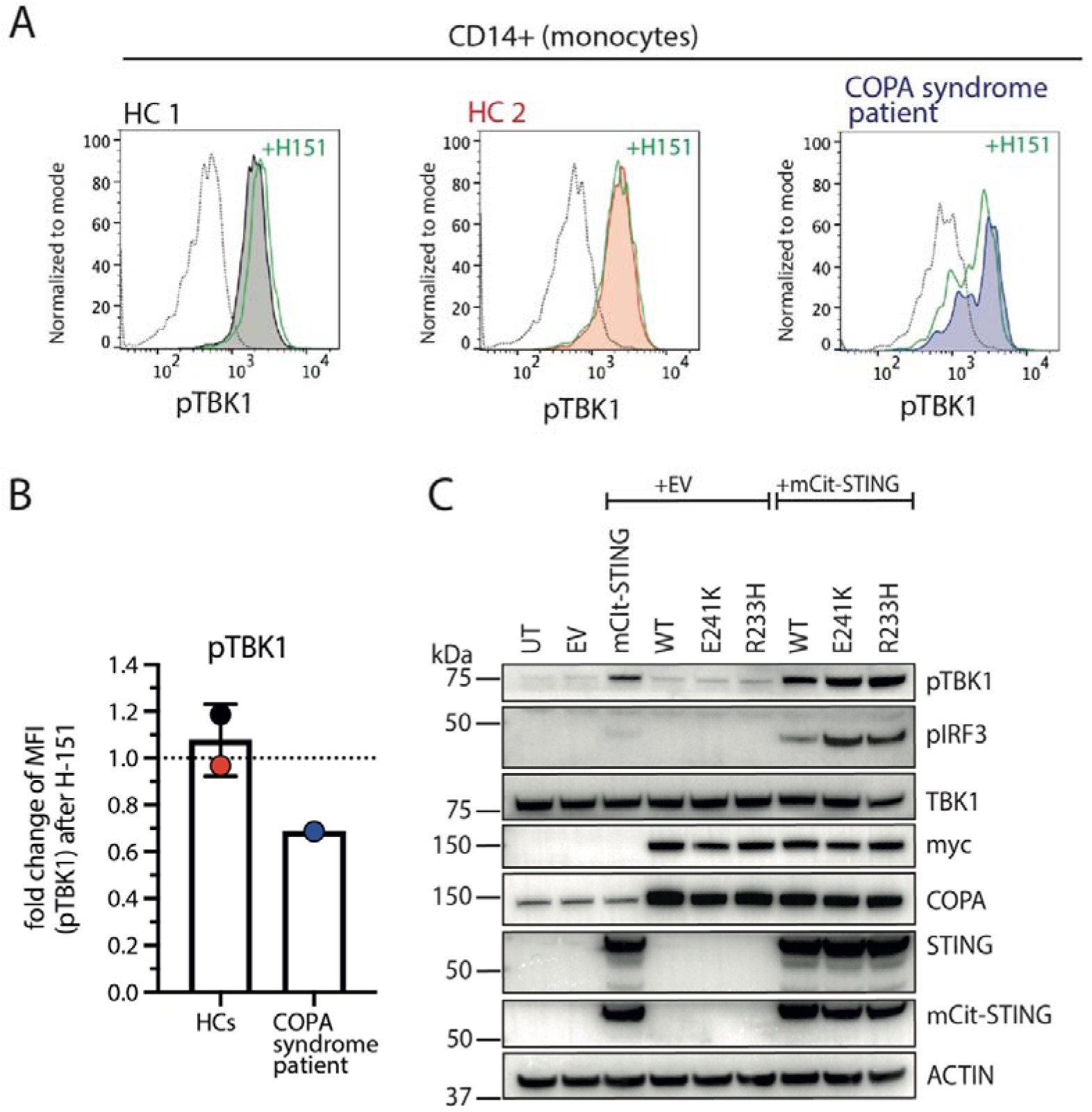
Mutations in COPA trigger STING-dependent inflammation. **A)** Representative histograms of flow cytometry analysis for phosphorylated TBK1 (pTBK1) of monocyte subpopulation (CD14+/CD3-) isolated from PBMCs of one COPA syndrome patient (blue) (3) and two healthy control (HC) individuals (black, red) before and after treatment with 5 μM of STING inhibitor H-151 for 4 hours (green line). Dotted line represents isotype control. n=1 **B)** Quantification of A). Bar graph represents fold change in pTBK1 mean fluorescence intensity (MFI) following H151 treatment relative to untreated control. Error bar represents standard deviation **C)** Analysis of inflammatory pathway activation in HEK293T cells following co-overexpression of STING and COPA mutants E241K and R233H (0.5μg DNA per construct). Immunoblots for phosphorylation of IRF3 (pIRF3) and TBK1 (pTBK1) were performed 24 hours after transfection. Representative experiment of at least 3 independent replicates is shown.

To further study inflammation in the context of COPA syndrome we generated overexpression plasmids encoding previously published loss-of-function COPA mutations E241K and R233H by mutagenesis PCR. We overexpressed these constructs in HEK293T cells which do not express detectable levels of endogenous STING (21, 39) and no basal activation of pTBK1 or pIRF3 was observed (Fig. 3C). Therefore, we co-expressed mCit-tagged STING together with the myc-tagged COPA mutant plasmids and could indeed detect increased levels of pIRF3 (Fig. 3C). Importantly, these experiments suggest that inflammation driven by COPA mutations in HEK239T cells only occurs in presence of STING, indicating the dysregulation of STING in COPA syndrome.

### Deficiency in COPI-mediated retrograde transport activates STING signalling

Besides COPA (subunit α), 6 other subunit proteins COP β, β’, δ, ε, γ, ξ as well as small GTPase Arf1 are essential for functional retrograde transport between Golgi and ER and within cis-Golgi compartments (Fig. 4A). The opposite transport direction between ER and Golgi (anterograde transport) is mediated by COPII complex subunits SEC13, SEC31, SEC23, SEC24 and small GTPase Sar1 (Fig. 4A). Here, we sought to determine whether activation of the STING signalling pathway is directly linked to loss of COPA function, or whether disruption of intracellular trafficking routes, both anterograde and retrograde, results in inflammation via this pathway. Therefore, we randomly selected COPI-subunit COPG1 and COPII-subunit SEC13 and used the CRISPR/Cas9 technology to delete these subunits in THP-1 cells.

**Figure 4:**
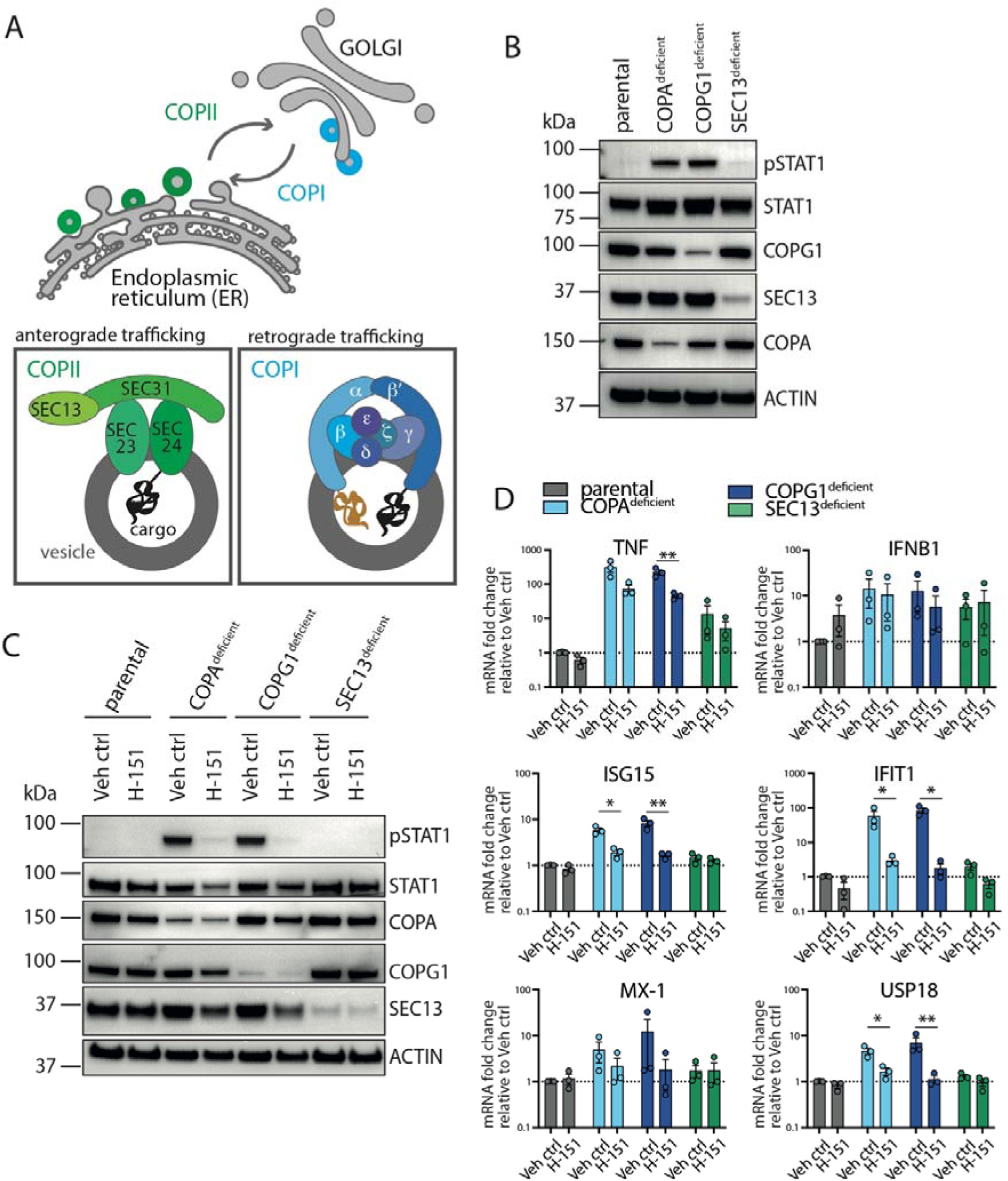
Deficiency in COPI-mediated retrograde trafficking activates the STING pathway. **A)** Schematic illustration of coatomer complexes COPII (green) and COPI (blue) mediating intracellular trafficking between endoplasmic reticulum (ER) and Golgi compartments. The COPII complex mediates anterograde transport from ER to Golgi. COPII vesicles formation (grey) at the ER involves inner subunits SEC23/24 and outer coat subunits SEC13/31. The COPI complex mediates retrograde transport from Golgi to ER as well as within cis-Golgi compartments. COPI vesicle formation occurs at the Golgi membrane through interaction of 7 subunits: α (COPA) and β’ (COPB1), β (COPB), δ (COPD), ε (COPE), γ (COPG), ζ (COPZ). After vesicle budding, the coatomer coats are shed off and released into the cytoplasm to allow vesicle fusion at the target membrane (2, 49). **B)** Immunoblot analysis shows CRISPR/Cas9-mediated genetic deletion of COPI-subunit protein COPG1 (encoding subunit γ) and COPII-subunit protein SEC13 in THP-1 cells. Baseline phosphorylation of STAT1 (pSTAT1) was assessed and is shown as a representative experiment of at least 3 independent replicates. **C)** COPG1^deficient^ and SEC13^deficient^ THP-1 monocytes were treated with STING inhibitor H-151 (2.5μM) for 11 hours and immunoblot analysis for pSTAT1 performed. Results are shown as a representative of 3 independent repeats. **D)** Results of qRT-PCR analysis of proinflammatory genes and ISGs in COPG1^deficient^ and SEC13^deficient^ THP-1 cells following H-151 treatment are shown. Data are pooled from 3 independent experiments and statistical significance was assessed by ratio-paired t-test. * p<0.05, ** p<0.01.

Interestingly, only deficiency in COPI complex proteins COPA and COPG1 but not the COPII protein SEC13 results in spontaneous phosphorylation of STAT1 and inflammatory gene transcription (Fig. 4B,D). In order to determine whether the inflammatory phenotype can be ameliorated by inhibition of STING signalling, THP-1 cells deficient for COPA, COPG1 or SEC13 were treated with STING inhibitor H-151. Immunoblot as well as qRT-PCR analysis show amelioration of pSTAT1 (Fig. 4C) and reduced transcription of proinflammatory genes and ISGs after inhibitor treatment (Fig. 4D). These findings indicate that indeed, the STING pathway is activated upon deletion of different COPI subunit proteins, however deletion of the COPII subunit protein SEC13 does not result in inflammation.

Collectively, our results demonstrate spontaneous activation of the STING signalling pathway in COPA syndrome. Furthermore, it appears disruption of COPI trafficking in general drives activation of STING signalling. Thus, mutations in other COPI proteins may underlie as yet undescribed disorders with immune/autoinflammatory pathology.

## Discussion

In this study, we have generated COPA syndrome model cell lines that recapitulate the IFN signature observed in PBMCs of COPA syndrome patient cells (3). We show that the type I IFN signature can be ablated by genetic deletion and pharmacological inhibition of STING. Interestingly, the clinical phenotype of COPA syndrome partially overlaps with symptoms reported in STING-associated vasculopathy with onset in infancy (SAVI), including severe systemic inflammation, recurrent fevers, interstitial lung disease, early onset in life and a predominant constitutive IFN gene activation (40). For SAVI, multiple case reports identified gain of function mutations in STING, causing spontaneous dimerization and therefore constitutive STING pathway activation (37, 41, 42). However, STING activation in COPA syndrome is less pronounced, as activated (phosphorylated) STING could not be detected in model cells lines and patient PBMCs (data not shown). Furthermore, we were not able to detect STING translocation to Golgi compartments in COPA^deficient^ HeLa cells, which is thought to be a prerequisite for STING signalling. However, COPA^deficient^ cells undergo partial Golgi fragmentation (38) similarly to cells treated with Brefeldin-A, a small GTPase inhibitor that prevents initiation of COPI and II vesicle formation and disrupts secretory trafficking (43). Therefore, STING signalling might occur off smaller Golgi fragments that could be below the level of detection, rather than from an intact Golgi complex, however this requires further investigation.

In our study we show that the STING pathway is activated upon deficiency of 2 different COPI-subunit proteins COPA and COPG1, but not COPII-subunit protein SEC13. Therefore, one could speculate that there may be other interferonopathies associated with defective retrograde transport caused by loss of COPI subunits other than COPA, that have not yet been described. Indeed, loss of function mutations in COPB2 (COPβ’) or COPD (COPδ, encoded by *ARCN1* gene) are linked to diseases associated with skeletal developmental defects and microcephalus formation (44, 45). Given the manifestation of these conditions, an underlying IFN signature may be present but undocumented. However numerous other factors would be contributing and determine whether a particular COPI-deficiency results in more pronounced developmental or inflammatory pathology. This becomes evident in COPA syndrome, which presents with incomplete penetrance leaving some mutation carriers unaffected. Co-factors contributing to disease onset in individuals with COPA mutation have not yet been identified but could be related to ER stress and the threshold for inflammatory signalling via cGAS/STING.

The finding that retrograde trafficking regulates STING is novel and the underlying mechanisms are not yet fully understood. Notable is that STING does not encode a dilysine motif (KKxx, KxKxx), which has been shown to be required for direct binding of COPA’s WD40 repeat domain to protein cargo (46). Therefore, retrograde transport of STING would require an adaptor protein. This hypothesis is supported by prepublication data from two laboratories, demonstrating that COPA binding to STING is dependent on Surfeit protein 4 (SURF4) (47, 48). This also agrees with our proteomic analysis, where SURF4 and COPA were immunoprecipitated with STING after overexpression in HEK293T cells (Figure 2A). Since STING signalling occurs from the ERGIC/Golgi compartments, it is conceivable that COPI-mediated retrograde transport may play an essential role to terminate signalling. However, whether activated STING is packaged into COPI vesicles and partially recycled to ER membranes or completely degraded via autophagy pathways remains to be determined. We were not able to visualize activated STING at the Golgi, likely due to the observed Golgi fragmentation following loss of COPA. The mechanism by which COPA deficiency leads to increased duration of activated STING signalling at the Golgi would also necessitate some initial signal for STING translocation from the ER. Whether homeostatic STING cycles between ER and Golgi or tonic cGAS activation is required to initially drive STING translocation in COPA syndrome, is currently under investigation.

Overall, this study shows that the type I IFN signature described in COPA syndrome patients is caused by a spontaneous activation of the STING pathway. This finding is particularly exciting in the context of pharmacological intervention. So far, COPA syndrome patients have been treated with various combinations of immunosuppressive drugs including DMARDs (disease-modifying anti-rheumatic drugs), NSAIDs (nonsteroidal anti-inflammatory drugs) and biologics, which were only able to partially control the disease (1, 3-10). Recently published case reports show treatment of two COPA syndrome patients with Janus kinase (JAK) 1/2 inhibitors Ruxolitinib or Barcitinib (10, 14). Although the IFN signature and joint inflammation were reduced successfully, patient lung function was not significantly improved (10, 14). Therefore, our study provides hope that targeting the STING pathway directly, for example with an inhibitor such as H-151, may be more specific and thus proves a greater beneficial approach to control type I IFN-mediated symptoms in COPA syndrome.

## Acknowledgement

The plasmids encoding mCitrine-fused to human STING, pEF-BOS empty vector control plasmid were kindly provided by V. Hornung (LMU Munich). We also thank C. Law and A. Shum (UCSF) for kindly providing the plasmid encoding COPA-myc-DDK. Single guide RNAs for MAVS and STING were kindly provided by Thomas Hayman (Melbourne University, WEHI) and Fiona Moghaddas (Melbourne University, WEHI), respectively. The HeLa cell line used in this study was kindly provided by C. Makhoul and P. Gleeson (Melbourne University). We thank Cynthia Louis (Melbourne University, WEHI) for providing human IL8 primers.

## Author Contributions

AS, SD, IP, KHS, CHY, DDN, LFD, CRH, PL and SV performed or assisted with experimentation. AS, SD, IP, KHS, CHY, DDN, LFD, CRH, PL, MPL, SV, MG and SLM were involved in experimental analysis and interpretation. All authors contributed to the writing of this manuscript.

## Abbreviations

COPI/II: coatomer complex I/II
COPA: coatomer complex I subunit A (α)
COPB: coatomer complex I subunit B (β)
COPE: coatomer complex I subunit E (ε)
COPD: coatomer complex I subunit D (δ)
COPG: coatomer complex I subunit G (γ)
COPZ: coatomer complex I subunit Z (ζ)
cGAS: cyclic GMP-AMP-synthase
CXCL10: CX-C Motif Chemokine Ligand 10
Dox: Doxycycline
ER: endoplasmic reticulum
GFP: green fluorescent protein
IFN: interferon
IRF3: Interferon regulatory factor 3
ISG15: interferon stimulated gene 15
JAK 1/2: Janus kinase 1/2
NF-κB: nuclear factor kappa
NLRP3: NOD-like receptor Pyrin Domain Containing 3
MAVS: mitochondrial antiviral signalling protein
MX1: MX Dynamin Like GTPase1
PBMC: peripheral blood mononuclear cell
PKR: protein kinase R
PRR: pattern recognition receptor
SEC13/31/23/24: COPII coat complex component
STAT1: signal transducer and activator of transcription 1
STING: stimulator of interferon gene
SURF4: Surfeit protein 4
TBK1: TANK binding kinase 1
TNF: tumor necrosis factor
UNC93B1: Unc-93 Homolog B1, TLR Signalling Regulator
USP18: Ubiquitin specific peptidase18

