## Supplementary Material for "Activation of STING due to COPI-deficiency"

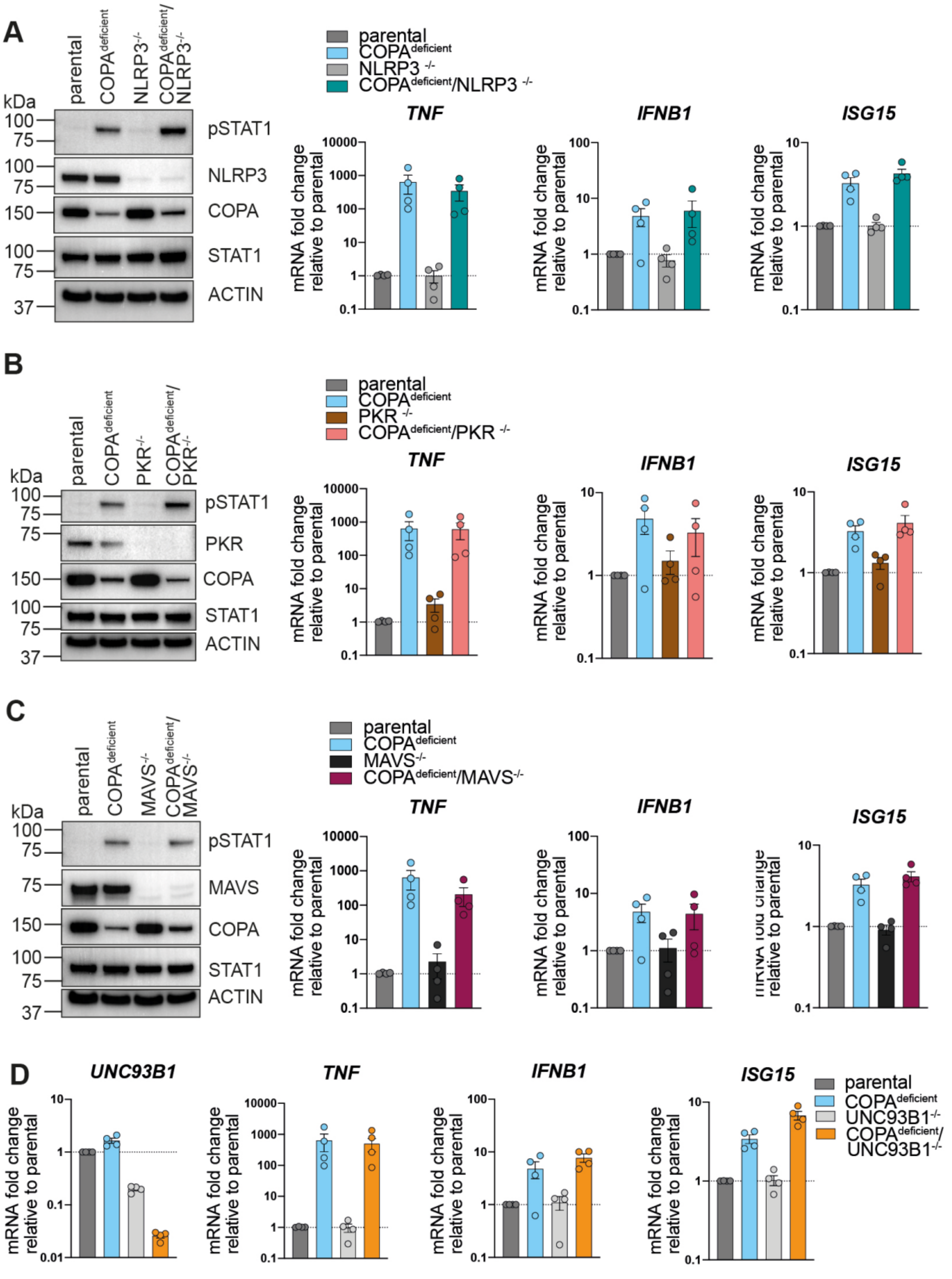


**Supplementary Figure 1. Candidate immune sensors that do not regulate inflammation associated with COPA-deficiency.**

The effect of genetic deletion of NLRP3 (A), MAVS (B), PKR (C), UNC93B1 (D) in COPA$^{deficient}$ THP-1 cells was assessed by immunoblot analysis of phosphorylated STAT1 (pSTAT1) and transcription analysis of *TNF*, *IFNB1* and *ISG15* by qRT-PCR. Data are means from 4 independent experiments. Error bars represent SEM.

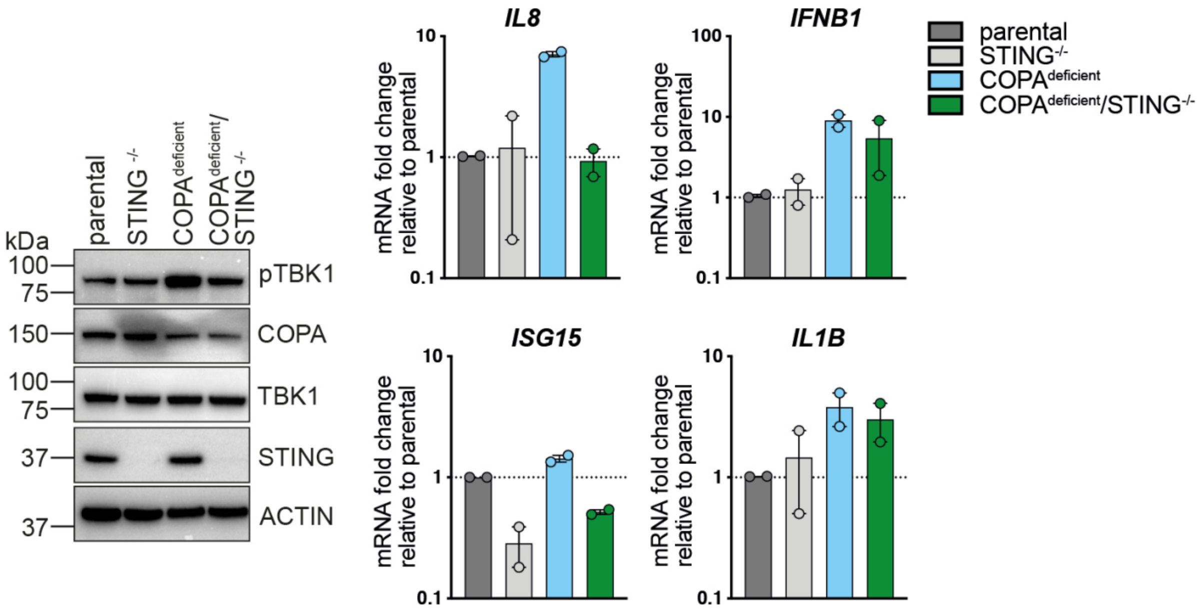


**Supplementary Figure 2. Inflammation in COPA$^{deficient}$ HeLa cells is STING-dependent.**

CRISPR/Cas9 gene editing technology was performed to genetically deleting STING in COPA$^{deficient}$ HeLa cells. Protein expression levels of COPA, STING and phosphorylated TBK1 (pTBK1) were assessed by immunoblot analysis of unstimulated cells (representative experiment shown). Inflammatory pathway activation was also investigated by qRT-PCR analysis of gene transcription profile of proinflammatory genes at baseline. Data were pooled from 2 independent experiments. Error bars represent SEM.

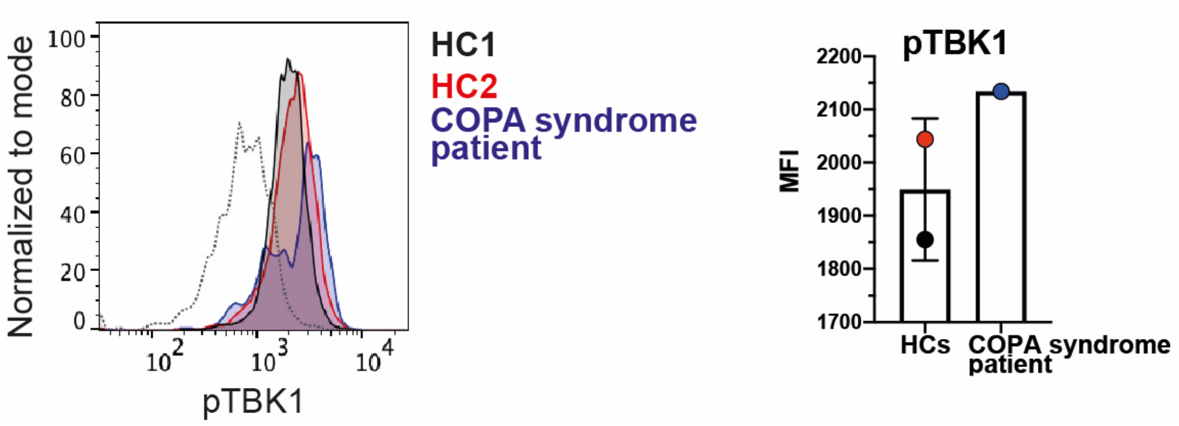


**Supplementary Figure 3. Phosphorylated TBK1 is elevated in monocyte population isolated from COPA syndrome patient PBMCs.**

Flow cytometry analysis of phosphorylated TBK1 (pTBK1) in monocytic subpopulation (CD14-positive, CD3-negative) of COPA syndrome patient PBMCs (blue) and 2 healthy individuals (HC in black and red) without further stimulation. Histogram shows data of a representative experiment, dotted line indicates isotype control. Column graph quantifies pTBK1 signal using mean fluorescence intensity (MFI) of the same experiment. Error bar represents standard deviation, n=1.

Supplementary Table 1: Single guide (sg) RNAs for CRISPR/Cas9-mediated gene editing.

| **Target gene** | **sgRNA sequence** |
|---|---|
| *COPA* (sgRNA1) | TAGATTGCCAGTTCCACACT |
| *COPA* (sgRNA2) | AATTCGAGACCAAGAGCGCG |
| *COPA* (sgRNA3) | ACATCCGATTCCACCGCACC |
| *STING* | AGAGCACACTCTCCGGTACC |
| *NLRP3* | TCCCGCTGGACCATCCTCGGCATG |
| *MAVS* | AGTACTTCATTGCGGCACTG |
| *PKR* | TAATACATACCGTCAGAAGC |
| *COPG1* | TGGTCAAGTAGCACATCCGA |
| *SEC13* | ATCCTTATCGCCGACCTCAG |
| *UNC93B1* | GGGCGTGCTCAAGAACGTGC |

Supplementary Table 2: Human primer sequences for quantitative Real-Time (qRT)-PCR for detection with SYBR Green.

| Target gene | F/R | Sequence (5’ → 3’) |
|---|---|---|
| *ACTIN* | F | GCGAGAAGATGACCCAGATC |
| | R | CCAGTGGTACGGCCAGAGG |
| *COPA* | F | ACTGGCAATCTAGAACCTGTG |
| | R | GACCAGAAATATCCCAAACGC |
| *TNF* | F | TCTCTCAGCTCCACGCCATT |
| | R | CCCAGGCAGTCAGATCATCTTC |
| *IFNA1* | F | GCCTCGCCCTTTGCTTTACT |
| | R | CTGTGGGTCTCAGGGAGATCA |
| *IFNB1* | F | TGTCGCCTACTACCTGTTGTGC |
| | R | AACTGCAACCTTTCGAAGCC |
| *ISG15* | F | TCCTGGTGAGGAATAACAAGGG |
| | R | GTCAGCCAGAACAGGTCGTC |
| *IFIT1* | F | ATCCACAAGACAGAATAGCCAG |
| | R | CCAGACTATCCTTGACCTGATG |
| *MX1* | F | GTTTCCGAAGTGGACATCGCA |
| | R | CTGCACAGGTTGTTCTCAGC |
| *USP18* | F | CCTGAGGCAAATCTGTCAGTC |
| | R | CGAACACCTGAATCAAGGAGTTA |
| *IL6* | F | TCAATATTAGAGTCTCAACCCCCA |
| | R | GAAGGCGCTTGTGGAGAAGG |
| *IL8* | F | CTGGCCGTGGCTCTCTTG |
| | R | CCTTGGCAAAACTGCACCTT |
| *IL1B* | F | AATCTGTACCTGTCCTGCGTGTT |
| | R | TGGGTAATTTTTGGGATCTACACTCT |
